# Real-time monitoring of endogenous Fgf8a gradient attests to its role as a morphogen during zebrafish gastrulation

**DOI:** 10.1101/2022.04.26.488902

**Authors:** Rohit Krishnan Harish, Mansi Gupta, Daniela Zöller, Hella Hartmann, Ali Gheisari, Anja Machate, Stefan Hans, Michael Brand

## Abstract

Morphogen gradients impart positional information to cells in a homogenous tissue field. Fgf8a, a highly conserved growth factor, has been proposed to act as a morphogen during zebrafish gastrulation. However, technical limitations have so far prevented direct visualization of the endogenous Fgf8a gradient and confirmation of its morphogenic activity. Here, we monitored Fgf8a propagation in the developing neural plate using a CRISPR/Cas9-mediated EGFP knock-in at the endogenous *fgf8a* locus. By combining sensitive imaging platforms with single-molecule Fluorescence Correlation Spectroscopy (FCS), we demonstrate that Fgf8a, produced at the embryonic margin, propagates by free diffusion through the extracellular space and forms a graded distribution towards the animal pole. Overlaying the Fgf8a gradient curve with expression profiles of its downstream targets determines the precise input-output relationship of Fgf8a mediated patterning. Manipulation of the Fgf8a input alters the signaling outcome, thereby establishing Fgf8a as a bona fide morphogen during zebrafish gastrulation. Furthermore, using diffusion-hindered versions of Fgf8a, we demonstrate that extracellular diffusion of the protein from the source is critical for it to achieve its morphogenic potential.

## Main text

The induction and organized arrangement of distinct cell types from a field of naïve cells is a fundamental challenge faced by all multicellular organisms during development. One mechanism to achieve this is to utilize secreted signaling molecules known as morphogens. These are produced from a localized source, distribute through the target tissue in a graded manner and elicit discrete cellular responses according to various concentration thresholds^1–4^. The Fibroblast Growth Factor 8a (Fgf8a) is one such molecule with inductive roles in mesoderm formation, neural patterning and organogenesis in vertebrates^5–8^. In the developing neural plate of early gastrulating zebrafish embryos, Fgf8a transcripts are detected near the embryonic margin and its target genes are expressed in increasingly broader domains away from the source^9,10^. Our previous study relying on mRNA micro-injections showed that fluorescently tagged Fgf8a, produced from ectopic sources in the embryo, is capable of forming a gradient of freely diffusing extracellular protein^11^. However, the distribution of Fgf8a, generated from its endogenous locus, has not been addressed due to the limitations in visualizing the endogenous protein. With recent advances in genome engineering^12–14^, it has now become possible to generate fluorescently tagged fusion constructs of Fgf8a from its endogenous locus and to monitor its propagation in real-time.

Hence, we first engineered an *fgf8a-EGFP* knock-in fish line by inserting the EGFP sequence into the endogenous *fgf8a* locus by CRISPR/Cas9^15,16^ (Supp fig. 1A). The resulting fusion protein has EGFP fused immediately after the signal peptide of Fgf8a, identical to the construct used in our earlier study^11^. *In situ* hybridization against *gfp* faithfully recapitulated the endogenous expression pattern of *fgf8a*, indicating that the fish line could be used as a valid read-out of the gene (Supp fig. 1B). The Fgf8a-EGFP fluorescence could be visualized in all prominent domains of *fgf8a* expression^6^- the dorsal marginal area at late gastrula, in the pre-somitic mesoderm as well as at the basal side of the midbrain-hindbrain boundary (MHB) during late somitogenesis stages (Supp fig. 1C). Homozygous *fgf8a-EGFP* embryos did not lack an MHB or a cerebellum and are viable as adults, as opposed to homozygous *acerebellar* (*ace*) loss of function mutant embryos, in which *fgf8a* is inactivated^6^, demonstrating the functionality of the *fgf8a* gene in the knock-in line (Supp fig. 1D).

Using conventional confocal microscopy, utilizing a photo-multiplier tube (PMT) for detection, it was not possible to visualize the endogenous Fgf8a-EGFP fluorescence before late gastrulation, owing to its weak expression levels at earlier stages. Therefore, we resorted to a sensitive quantitative imaging protocol, using GaAsP hybrid detectors and spectral auto-fluorescence subtraction^17, 18^ (see Methods, Suppl fig. 2A). This enabled us to visualize the endogenous Fgf8a-EGFP fluorescence in the neural plate of early and mid-gastrulating transgenic embryos (Fig. 1A; Suppl fig. 2B). By analysing the signal intensity in the neural plate along the animal-vegetal axis, we found that Fgf8a-EGFP, secreted from its source at the embryonic margin, forms a graded distribution towards the animal pole (Fig. 1B; Suppl fig. 2B). Additionally, we also examined the Fgf8a-EGFP fluorescence intensity near the margin, along the dorsal-ventral axis, at mid-gastrula. This showed that during gastrulation, Fgf8a-EGFP does not only form a gradient along the animal-vegetal, but also along the dorsal-ventral axis (Suppl fig. 2C). While the former is generated by propagation of the protein from a localized source, the latter could be a direct manifestation of the dorsal to ventral mRNA gradient of *fgf8a*, which has been previously suggested to regulate dorsoventral patterning of the embryo^19,6^.

**Figure 1.**
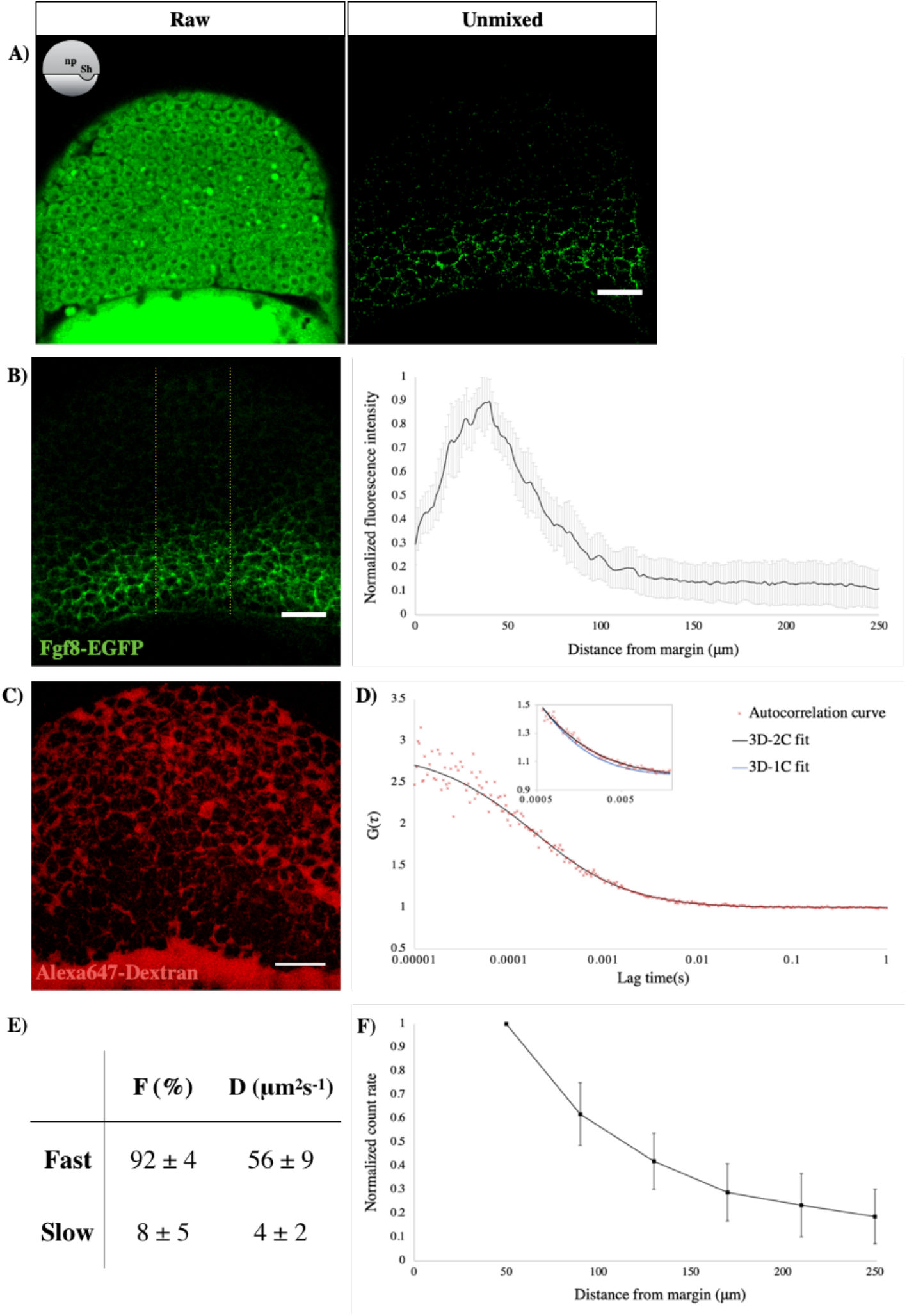
Fgf8a-EGFP forms a gradient during gastrulation by freely diffusing via extracellular spaces. **A)** EGFP fluorescence in an optical section of the neural plate in early gastrula-staged *fgf8a-EGFP* embryos, as visualised using the GaAsP detector, before (left) and after (right) linear un-mixing. The orientation of embryos shown as scheme. np-neural plate, Sh-shield. **B)** Sum-intensity projection of un-mixed images along z-axis (left) used to extract fluorescence intensity profiles along a vertical area (dotted yellow bounds). The analysis reveals a graded distribution of Fgf8a in the animal-vegetal axis (right). N=15. **C)** Extracellular spaces (ECS) in an early gastrula embryo visualised by Alexa-647 tagged Dextran injection. **D)** FCS autocorrelation curve from ECS of *Tg(fgf8a:fgf8a-EGFP)* embryo fitted with 2-C (black curve) and 1-C (inset-blue curve) 3D-diffusion models. **E)** Results from fitting with 2C-3D diffusion model. F, fraction of each component and D, diffusion coefficient. n=63. **F)** Plot of count rate, derived from FCS analysis, vs distance from margin in early gastrula embryos showing graded distribution of Fgf8a in the ECS along the animal-vegetal axis. N=20. All scale bars-50μm.

Having visualized the animal-vegetal gradient of endogenous Fgf8a-EGFP, we then aimed to understand how the molecules, produced at the embryonic margin, traversed towards the animal pole. While several mechanisms of morphogen transport have been identified, the most prevalent mode of transport is extracellular diffusion. Our earlier study with exogenous Fgf8a-EGFP showed that it propagates primarily by random diffusion via the extracellular space^11^. However, there could be several artifacts associated with the ectopic expression of a protein, such as aggregation or mislocalization, due to which it was important to examine the endogenous species. To this end, we employed the single-molecule analysis technique of FCS^11,20,21^ in the extracellular space of *fgf8a-EGFP* transgenics at early gastrula stages. The extracellular space was specifically labelled by injecting a non-cell permeable tracer, Alexa647-tagged dextran (Fig. 1C). All FCS auto-correlation curves for extracellular Fgf8a-EGFP could be fitted with a 3-dimensional (3D) random diffusion model. Although some of the curves fitted with a 1-component 3D diffusion model (3D-1C), in which only a single species of molecules is assumed, the majority of the curves could not be described by this model within the lag times of 0.5-50 ms (Fig. 1D-inset). Instead, these curves fitted well with a 2-component 3D diffusion model (3D-2C), suggesting the presence of two species of molecules- a major proportion (92%), moving with a diffusion coefficient (D) of 56 μm^2^s^-1^, and a remaining slow moving fraction (D = 4 μm^2^s^-1^) (Fig. 1D, E). The value of D_fast_ is in the same order of magnitude as that of monomeric EGFP in solution^22^, hence reflecting free diffusion. In order to further demonstrate that the endogenous Fgf8a-EGFP forms a gradient by extracellular dispersal, we recorded the FCS fluorescence count rates, as a direct measure of the molecular concentration of the protein^11^, from the extracellular space within the neural plate, at varying positions along the animal-vegetal axis. This showed a smooth gradual decrease in protein availability with increasing distances from the embryonic margin (Fig. 1F).

Next, we sought to decipher how the endogenous Fgf8a-EGFP gradient correlates with discrete domains and levels of target gene expression in the gastrulating embryo, which would lend an additional descriptive support to the morphogenic action of Fgf8a. To achieve this, we first performed single molecule fluorescence *in situ* hybridization ^23^ against the Fgf8a target genes^24–28^ *tbxta* and *etv4*, in whole mount early gastrula-staged embryos. While the *tbxta* foci were specifically detected only within 12 cell tiers from the embryonic margin, there were no locations devoid of *etv4* transcripts (Fig. 2A,C). To confirm the specificity of the *etv4* probe set, we inhibited Fgf signaling by treating embryos, from pre-blastula stages, with 10 μM SU5402. No *etv4* foci could now be detected in such embryos at early gastrula, in contrast with the DMSO-treated controls (Fig. 2B). Thereafter, we quantified the expression profiles of both *tbxta* and *etv4*, along the animal-vegetal axis, in the developing neural plate, by counting the number of foci, and plotting it as a function of distance from the embryonic margin. This was further overlaid with the Fgf8a gradient data, as obtained using FCS, to determine the various thresholds of the signaling molecule required for the induction of distinct cellular responses. The *tbxta* profile was found to closely follow the Fgf8a gradient curve, with the maximum number of foci within 40-70 μm from the margin, corresponding to an Fgf8a availability of at least 80 percent of its source concentration (0.8*C_0_). Thereafter, the number of foci decreased steeply, before nullifying at 0.4*C_0_ of Fgf8a. For *etv4*, while the transcripts were detected all the way up to the animal pole, its peak expression domain was identified to be shifted when compared to that of *tbxta*, with the highest number of *etv4*-positive foci detected within 70 and 140 μm from the embryonic margin. This correlated with extracellular Fgf8a levels of 0.8*C_0_ and 0.4*C_0_, respectively (Fig. 2C). In addition to the single molecule fluorescence *in situ* hybridization approach, we also utilized a *Tg(etv5b:etv5b-Venus)* knock-in reporter line to quantitate the expression profile of Etv5b-Venus another Fgf8a target gene^27,28^. The transgenic line was generated via CRISPR/Cas9-mediated homologous recombination, by which the Venus fluorophore sequence was inserted right before the start codon of the endogenous *etv5b* locus. *In situ* hybridization against *etv5b* and *venus* showed identical localization pattern of the transcripts and the fusion protein was found to be localized to the nucleus, consistent with the role of Etv5b as a transcription factor (Suppl fig. 3). This nuclear signal was completely lost upon incubation with 10 μM SU5402, demonstrating the validity of the transgenic line (Fig. 2D). By measuring the nuclear fluorescence intensity at increasing distances from the embryonic margin, we found a very shallow gradation in protein abundance across the animal-vegetal axis, with peak intensity up to 140 μm from the embryonic margin, where ~0.4*C_0_ of Fgf8a-EGFP is likely available for signaling (Fig. 2E).

**Figure 2.**
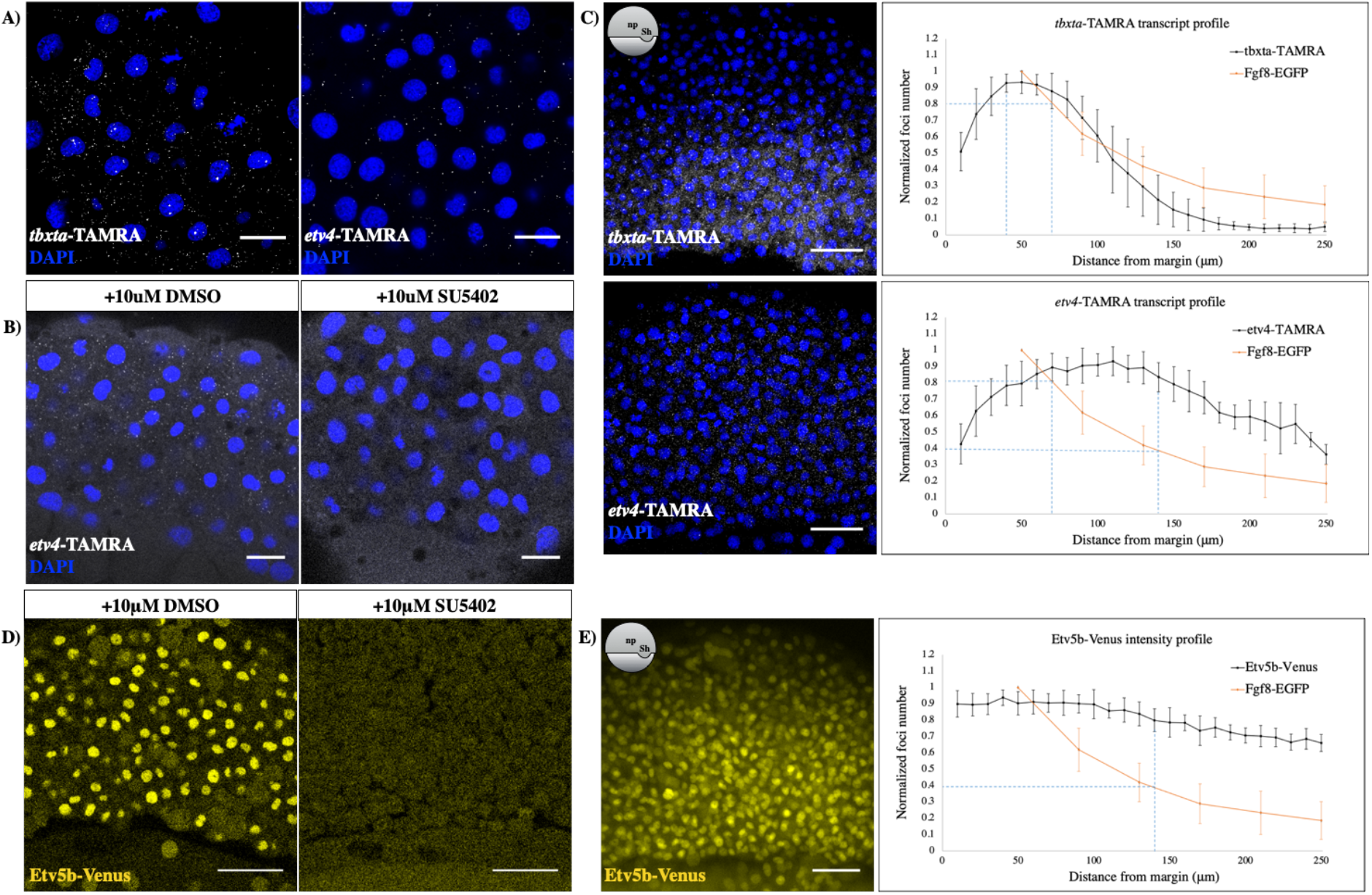
Recording the expression profiles of Fgf8a target genes during gastrulation. **A)** Optical sections from smFISH against *tbxta* and *etv4* in early-gastrula staged embryos, after threshold adjustment for background subtraction. **B)** smFISH against *etv4* in DMSO (left) and SU5402 (right) treated embryos at early gastrulation. Optical sections are shown. Scale bars for A & B-20μm; lateral views are shown. **C)** Maximum-intensity z-projected images for smFISH against *tbxta* (top) and *etv4* (bottom) in early gastrula-staged embryos (left) and plot of normalised transcript number vs distance from margin (right). N=10. For A-C, DAPI (blue) used as nuclear marker. Overlay with the Fgf8a distribution profile (orange curve) shows its extracellular concentration corresponding to peak expression levels of both the transcripts (see blue dotted lines). **D)** Venus fluorescence in DMSO (left) and SU5402 (right) treated *Tg(etv5b:etv5b-Venus)* embryos at early gastrulation. Optical sections of laterally mounted embryos are shown. **E)** Sum-intensity z-projected image of Etv5b-Venus from early gastrula-staged transgenic embryos (left) and analysis of nuclear fluorescence intensity as a function of distance from the margin (right). Overlay with the Fgf8a gradient curve (orange curve) shows its minimum relative abundance in the extracellular spaces that correspond to the peak Etv5b output (see blue dotted lines) N=10. Scale bars for C to E-50μm. For C & E, exact orientation is shown as schematics.

Our FCS experiments with the endogenous Fgf8a-EGFP earlier showed that, while a major fraction of molecules disperse by free diffusion, a minor proportion remains relatively immobile (see Fig. 1E). Based on previous studies^11,29^, we reasoned that this slow-moving fraction results from interaction with the extracellular matrix constituents, particularly heparan sulphate proteoglycans (HSPGs). In accordance with this, injection of Heparinase I (), an enzyme that cleaves heparan sulphate side chains of HSPGs^30^, into the *fgf8a-EGFP* transgenics, diminished the slow-moving fraction significantly (Fig. 3A). However, HepI injections also resulted in an overall increase in the Fgf8a-EGFP levels in the extracellular space, possibly due to the disassociation of Fgf8a from cell surface HSPGs (Fig. 3B). Thereafter, we examined the effect of HepI injections on the Fgf8a extracellular concentration profile in the neural plate at early gastrula, by analysing the FCS fluorescence count rates at increasing distances from the embryonic margin. The application of HepI was found to result in a shallower gradient (Fig. 3C), which not only emphasized the role of Fgf8a-HSPG interactions in shaping the protein distribution, but also provided us with a technique to manipulate the extracellular thresholds of ligand input. In order to determine whether this change in Fgf8a-EGFP input leads to a concomitant change in the range of signaling outcomes, we first performed *in situ* hybridization against the Fgf8a target genes^31^ *spry2* and *tbxta* on early gastrula-staged control and HepI-injected embryos. The expression domains of both target genes were found to broaden towards the animal pole upon HepI injection (Fig. 3D). To further quantitate the effect of HepI-mediated manipulation of the Fgf8a distribution, we used single molecule fluorescence *in situ* hybridization against *tbxta* in HepI-injected embryos at early gastrula stage and analysed its foci number as a function of distance from the embryonic margin. The amplitude of its expression profile was found to decrease when compared to the control embryos, possibly due to the cleavage of HSPG side chains from the cell surface, which affects the ligand-receptor interaction and begets a reduction in signaling strength. Nevertheless, the *tbxta* expression domain itself was found to increase by 2-3 cell tiers, on average, upon HepI injection (Fig. 3E). Thus, flattening the Fgf8a extracellular gradient alters the domains of induction of its downstream targets, providing a functional proof for the morphogenic action of endogenous Fgf8a, in accordance with Lewis Wolpert’s French flag hypothesis.

**Figure 3.**
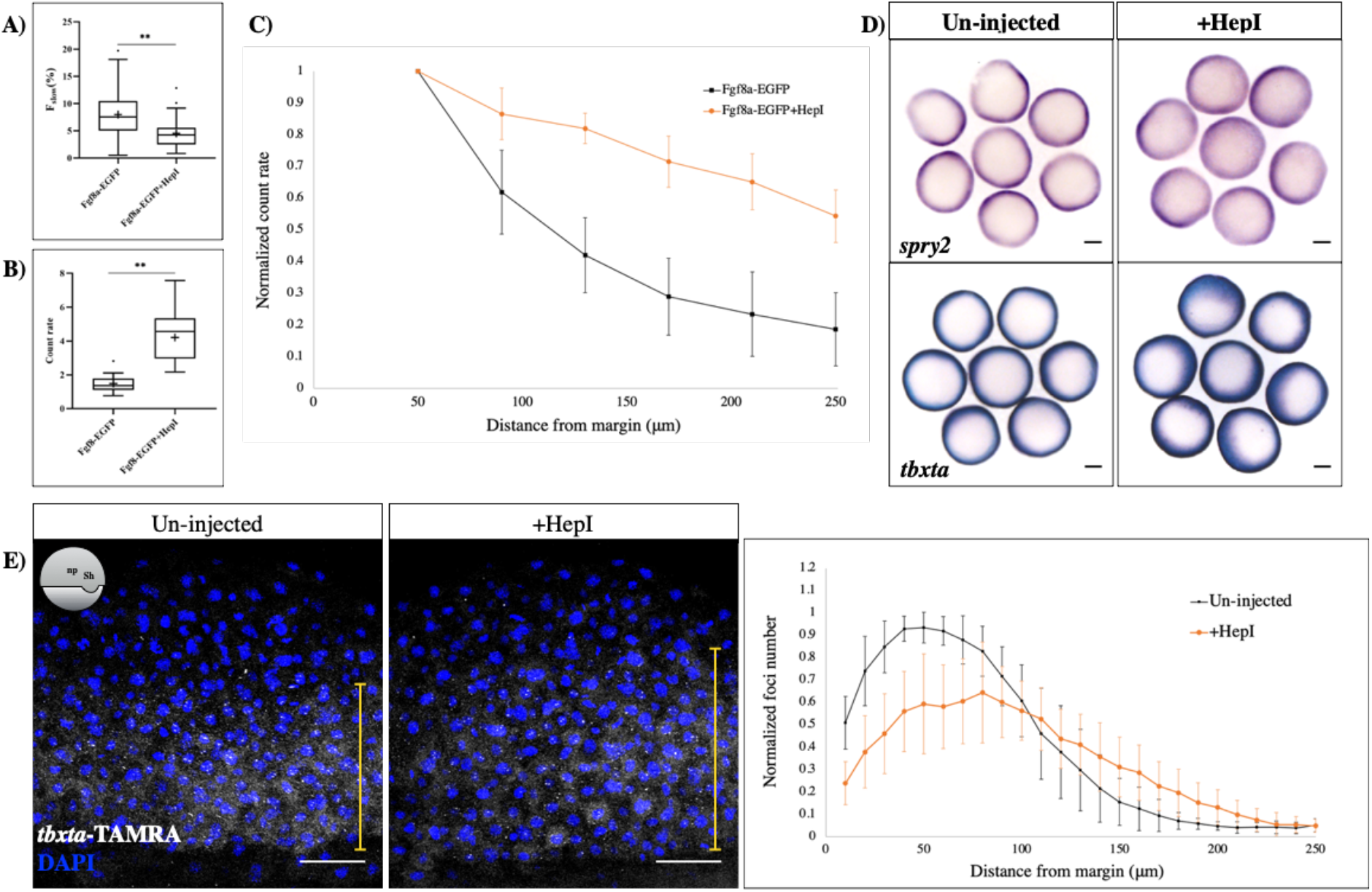
Manipulation of Fgf8a gradient by HepI injection alters the signalling output. **A)** Injection of HepI reduces the proportion of slow moving fraction of Fgf8a, as detected by FCS, in the extracellular spaces. Mean ± s.d. is shown. n=63 for un-injected, n=56 for HepI. Student’s t-test P value = 1.2* 10^-7^. **B)** Box plot showing range of Fgf8a-EGFP count rates in extracellular spaces at the embryonic margin in un-injected and HepI injected embryos. Note the increase in count rates upon HepI injection. n=20. ‘x’ denotes the mean. Student’s t-test P value = 4.5*10^-7^. **C)** Plot of count rate vs distance from margin in control (black) and HepI (orange) injected cases. N=20 for control, N=13 for HepI. **D)** *In situ* hybridisation against *spry2* (top) & *tbxta* (bottom) in control and HepI injected early gastrula embryos. Animal pole views are shown. Scale bars-100μm. **E)** smFISH against tbxta in un-injected and HepI-injected early gastrula staged embryos and plot of normalised transcript number vs distance from the margin. Note the reduction in *tbxta* expression near the margin and increase in its range of induction away from the margin (range depicted by the yellow line over the images), upon HepI injection. Orientation of embryos shown as scheme. N=10. Scale bars-50μm.

Finally, we aimed to test the importance of extracellular diffusion in long-range dispersal and morphogenic activity of Fgf8a. While extracellular diffusion has been at the forefront of the transport models for morphogen gradient formation, the reliability of such diffusion-based gradients in accurately positioning the boundaries of distinct target cell responses and their reproducibility across individuals are not fully understood. Also, it has been a matter of debate whether relying primarily on a passive process such as random-walk diffusion can generate suitable gradients within the necessary time frame across a sufficiently large field. Several alternatives to extracellular diffusion have been proposed, among which, directed transport by specialized cellular extensions termed cytonemes has garnered much attention and has also been reported for Fgfs in Drosophila. Therefore, we engineered two diffusion-abrogated versions of Fgf8a^32^. EGFP-labelled Fgf8a was attached either to the transmembrane domain of human T-cell surface protein Cluster of differentiation-4 (CD4), or the zebrafish full-length transmembrane protein ß-Sarcoglycan (Sgcb). Un-tethered Fgf8a-EGFP, as in our earlier study^11^, was used for comparison (Fig. 4A). The respective constructs were micro-injected as mRNA into 1 cell-staged embryos and imaged at mid-blastula stages. While the un-tethered Fgf8a-EGFP was abundantly secreted into the extracellular space, the CD4-tagged version was primarily anchored to the cell membrane, and the Sgcb-linked protein was located both intracellularly and in patches at the cell membrane (Fig. 4B). Thereafter, we generated ectopic clones expressing the respective fusion proteins in the developing embryo, A membrane marker, hRas-mKate, was co-injected to precisely label the source cells, and imaging was done at mid-blastula stages to determine the dispersal of the proteins from their respective sources. In control embryos, Fgf8a-EGFP diffused away from its source, via the extracellular space, and filled the entire animal pole of the embryo. In stark contrast, CD4-Fgf8a-EGFP could be found only in cells up to 3-4 cell diameters away from its source while Sgcb-Fgf8a-EGFP was primarily retained within the source cells (Fig. 4C). Interestingly, in Sgcb-Fgf8a-EGFP-injected embryos, some protein clusters could be detected at the tips of mKate-labelled membranous protrusions arising from the source cells (Suppl fig. 4). The CD4-tethered version could also be using this mechanism to traverse to the adjacent cell surfaces, although we have not been able to monitor this. Such membranous protrusions have earlier been shown to be utilized in the transport of Wnt8a in the zebrafish developing neural plate. While it is endogenous lipidation that confers Wnt8a its ability to localize to the membrane tips, it is possibly the engineered tethers in our study. Subsequently, we performed *in situ* hybridization against the Fgf8a target gene^33^ *spry4* on such embryos at late blastula stages. An additional probe against *gfp* was utilized to precisely detect the source cells. This showed that while the free diffusing Fgf8a-EGFP induces *spry4*expression at a distance surrounding the source cells, CD4-Fgf8a-EGFP could activate it only in the nearest neighbours and Sgcb-Fgf8a-EGFP was completely incapable of *spry4* activation outside the source cells (Fig. 4D, E). Thus, we conclude that extracellular diffusion is the predominant mechanism by which Fgf8a propagates over a long range and induces downstream signaling events, thereby ensuring proper patterning of the tissue.

**Figure 4.**
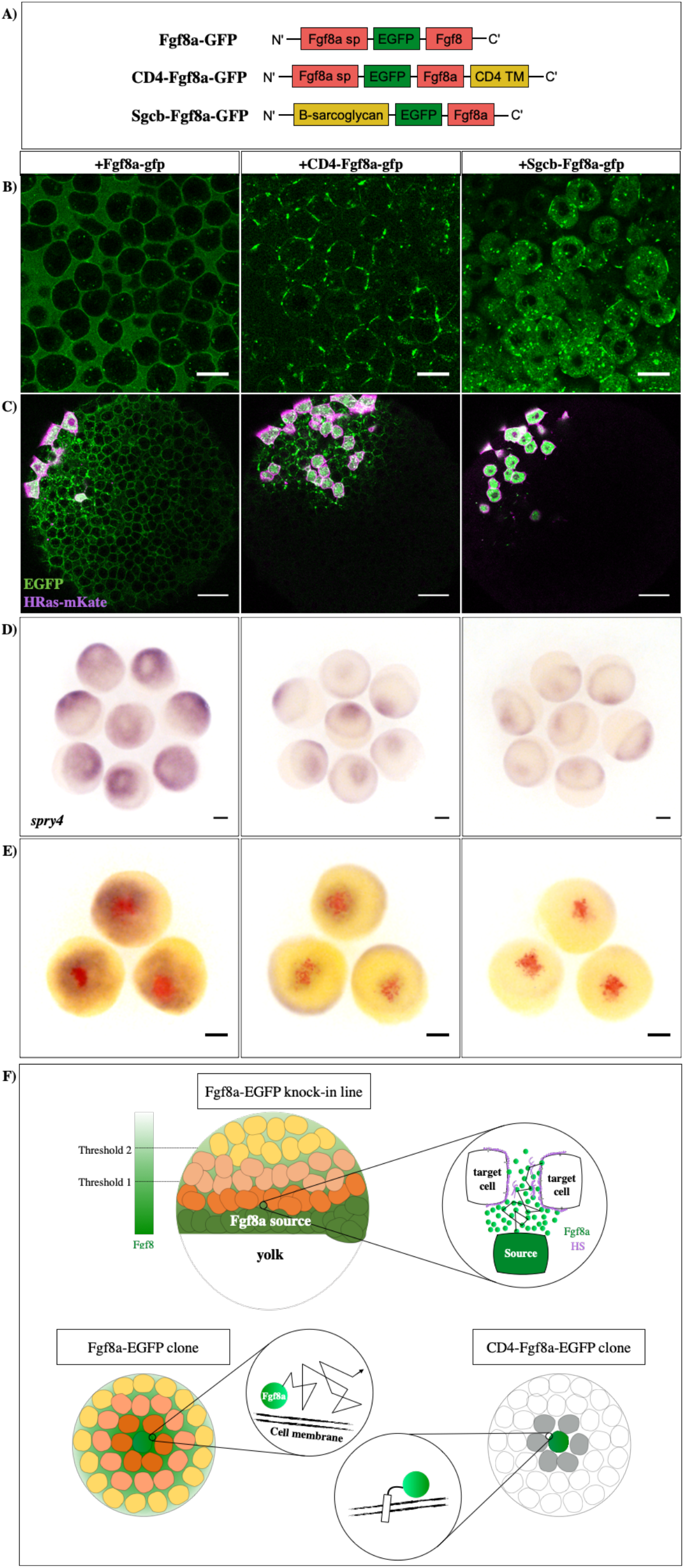
Diffusion of Fgf8a is necessary for its morphogenic activity. **A)** The three different versions of Fgf8a used in this study. sp-signal peptide. **B)** Micro-injection of the constructs at 1-cell stage and imaging at blastula stages to assess localisation. Scale bars-20μm. **C)** Generation of ectopic clones of the constructs and imaging at late-blastula stages to determine protein spreading. Co-injection with HRas-mKate to label the source cells (magenta). Scale bars-50μm. **D)***In situ* hybridisation against *spry4* at late blastula after generating ectopic clones for all versions of Fgf8a. **E)** Double *in situ* hybridisation against *gfp* (red) and *spry4* (blue) in such embryos at late blastula stage to depict the ectopic source and target gene induction domain respectively. For both D & E, scale bars-100μm. Animal pole views of embryos shown in all panels. **F)** Summary of the work: Visualising endogenous Fgf8a via knock-in reveals its graded distribution in extracellular space (Upper panel, left). Single molecule FCS reveals Fgf8a as a freely-diffusing morphogen (enlarge on right). **B)** Membrane-tethering of Fgf8a in cell clones restricts its signalling range (lower panel).

Overall, making use of the technical advancements in gene editing and signal detection, we have been able to follow in real-time, the distribution of endogenous Fgf8a molecules during zebrafish gastrulation. We find that Fgf8a functions as a *bona fide* morphogen in the developing neural plate, by establishing a concentration gradient and eliciting threshold dependent cellular responses, in support of Lewis Wolpert’s French flag hypothesis^1^. We also provide direct evidence that a passive process, such as free extracellular diffusion, is sufficient for a morphogen to disperse across its target tissue within the necessary time frame. Although other studies have previously implicated cytonemes as an alternate means of Fgf ligand delivery^34,35^, we show here that it is critical for the ligand to disperse freely via the extracellular space to form a long-range gradient (Fig. 4F). Future work will aim to hinder the diffusion of endogenous Fgf8a and evaluate its repercussions on normal patterning events during gastrulation. Additionally, it will be key to understand the distribution dynamics of other Fgfs, co-expressed with Fgf8a during gastrulation, to fully comprehend the intricate network which is in place to guide the development of a multicellular organism with precision.

## Methods

### Zebrafish husbandry

The fishes were raised and housed in the facilities of the Biotechnology Center of TU Dresden (BIOTEC) and Center for Regenerative Therapies, Dresden (CRTD) respectively, as per conditions described elsewhere^37^. The adult fish of the required genotype were allowed to mate in specially designed containers filled with fish water. Embryos were collected in petri-dishes filled with E3 medium and stored mostly at 28°C. The staging was done according to standard criteria^38^. Live embryos were kept for a maximum of 4 days before discarding or handing over to the facility for raising. AB fish were used as the wildtype (WT AB) in this study.

All animal experiments were carried out in accordance with animal welfare laws of the Federal Republic of Germany (Tierschutzgesetz) that were enforced and approved by the competent local authority (Landesdirektion Sachsen; protocol numbers TVV21/2018; DD24-5131/346/11 and DD24-5131/346/12).

### CRISPR/Cas9 transgenesis

For the generation of *fgf8a-EGFP* transgenics, two sgRNA target sites (ts1: GGTGAAGGAATATTTAAAAG<u>TGG</u>; ts2: GGAGAGGAGGGAACGTCTTC<u>AGG</u>; PAM sites are underlined) were chosen as shown in Suppl fig 1A. sgRNAs and Cas9 mRNA were synthesised as described elsewhere^39^. The bait sequence, as illustrated in Suppl fig 1A, was cloned into pCS2+ vector. 20 pg of this plasmid was injected along with 50 pg of both sgRNAs and 150 pg of Cas9 mRNA, into 1-cell staged wildtype (AB) embryos. F0 adults were screened for germ-line insertion by outcrossing to wildtype, extracting the genomic DNA from their progeny, and amplification by PCR, with one primer targeted to EGFP and the other outside the bait (Fw: GCCTGGCAAGAAATGGGACA; Rv: CAGGGTCAGCTTGCCGTAG). The transgenic line was maintained by outcrossing to wildtype. To verify single integration, PCR amplification was carried out from the genomic DNA of animals homozygous for the transgene, with primers annealing outside the homology arms of the bait (Fw: GCCTGGCAAGAAATGGGACA; Rv: CTCGACTCCCAAATGTGTCCGT), which revealed only one band of the expected size. Sequencing of this amplicon revealed no changes other than a 6bp deletion in the intronic sequence corresponding to ts2.

For the generation of *etv5b-venus* transgenics, a single sgRNA target site (ts: GGAACGCACAGGGATAAACG<u>GGG</u>), with minimal probability of off-target effects according to CRISPRscan.org^40^, was utilized (Suppl fig 3). The aforementioned protocol was repeated to execute the insertion. F0 adults were screened for germ-line insertion by PCR amplification from the genomic DNA of progeny, with one primer targeted to Venus and the other outside the bait sequence (Fw: CAGAGGATTATACAAACGCTGGG; Rv: ACTTGTGGCCGTTTACGTCG).

### Micro-injections

During mRNA micro-injections, 100 pg of the constructs were usually injected either into the cytoplasm of 1-cell staged embryos, or into a single cell of 32-cell staged embryos. For comparison of the diffusion-hindered Fgf8a versions, 100 pg of Fgf8a-EGFP mRNA was injected as control, and equimolar amounts of the other constructs were injected. For visualization of the extracellular space, 0.2 nl of 5 μM solution of Alexa fluor 647-labelled Dextran (Invitrogen) was injected into the animal pole of sphere-staged embryos. In the case of Heparinase I (Sigma-Aldrich), 0.2 nl of 1unit/μl solution of Heparinase I was injected 5-6 times at various positions of sphere-staged embryos to ensure an even distribution of the enzyme.

### SU5402 treatments

Embryos were manually dechorionated and treated from pre-blastula stages with 10 μM SU5402 or DMSO, and fixed or imaged at early gastrula.

### *In situ* hybridization

Whole-mount *in situ* hybridization was carried out as described previously^6^. Probes were prepared from linearized plasmid templates using the DIG/Flu RNA labelling mix (Roche). 1:4000 of anti-dig-AP and 1:1000 of anti-flu-AP antibodies (Roche) were utilized for the staining procedure. BM purple (Roche) and FastRed substrate (Sigma-Aldrich) were used to develop the blue and red staining, respectively.

A previously established protocol^23^ was used to perform smFISH. While oligos against *etv4* were designed using the Stellaris Probe Designer, a known probe set^23^ was utilized for *tbxta*. TAMRA-labelled (at 3’ end) probe sets for both genes were synthesised from Biosearch technologies. A final probe concentration of 250 nM was utilized for detection.

### Imaging

Live or fixed embryos were appropriately mounted and oriented either in 1% low-melting agarose in E3 or 70 % glycerol (whole-mount *in situ* samples). All stereo-microscope derived images were taken using an Olympus MVX10 MacroView fluorescence microscope. Confocal images were acquired using a 40X C-Apochromat NA1.2 water immersion objective of the Zeiss LSM780/Confocor3 microscope, at a room temperature of 22.5 °C.

### GaAsP imaging protocol

For visualisation of endogenous Fgf8a-EGFP, confocal imaging was implemented using the GaAsP hybrid detector^17^. This allowed for simultaneous acquisition of spectral information from the samples, which was coupled to the mathematical technique of linear un-mixing using ZEN (Zeiss), to separate the various spectral profiles emanating from the sample^18^. *Tg(bactin:hRas-EGFP)* embryos, in which EGFP is found in abundance, and gastrula-staged wildtype embryos, were imaged using the GaAsP, to obtain the specific emission spectrum of EGFP and various spectra corresponding to background autofluorescence, respectively. Such profiles were subsequently used as references for linear un-mixing of images derived from *fgf8a-EGFP* to subtract the pixels corresponding to background autofluorescence and visualize only those specific to EGFP.

### Fluorescence correlation spectroscopy (FCS)

FCS measurements were performed using an in-built FCS setup of the Zeiss LSM780/Confocor3 microscope, with the 40X C-Apochromat NA1.2 water immersion objective, at a room temperature of 22.5 °C. A 488 nm Argon laser (laser power-15 μW) was utilized for excitation and a BP 505-540 IR emission filter was used for detection, with the pinhole set to 1AU. On each day of measurement, the microscope was calibrated for the correction collar and pinhole alignment using 50 μl of 15 nM Alexa fluor 488 (A488). For the experimental samples, each position of measurement was manually selected by acquiring a confocal image (of A647-Dextran) using the appropriate setting and 5 repetitions of 10 s-long FCS measurements were taken.

Subsequent analysis was done as described previously^11^. In short, the curves obtained for A488 were fit with a 3-dimensional 1-component diffusion model (3D-1C) using ZEN (Zeiss) and the average dwell time (τ_D_) for the dye, in the observation volume, was determined. The triplet fraction and triplet relaxation time were fixed at 10 % and 30 μs, respectively, while the structural parameter was allowed to fit freely. The average value obtained for τD was then used to calculate the lateral radius (ω) of detection volume, using the equation D =ω^2^/4τ_D_, where D is the diffusion coefficient (literature value) of A488 in solution. For all experimental samples, the FCS auto-correlation curves were fit with either 3D-1C or 3D-2C diffusion models and the efficiency of fitting was compared. Again, the triplet fraction and triplet relaxation time were fixed at 10 % and 30 μs, respectively, while the structural parameter was fixed as the average obtained for A488 (~5). The dwell times obtained from the appropriate fit and the lateral radius of detection volume, determined earlier, were used to calculate the diffusion coefficients of the molecules.

### Fluorescence intensity analysis

Confocal images were analyzed for overall fluorescence intensity using Fiji/ImageJ. Optical sections for a given sample were sum-intensity projected along the z-axis and the intensity profile across a 100 μm wide region, corresponding to the neural plate, was extracted as a function of the distance from the embryonic margin. For each embryo, the profile was normalized using the maximum intensity value within that sample.

For analysis of nuclear fluorescence intensity, confocal z-stacks were processed using a denoising filter, followed by nuclear segmentation using a blob-finder analysis operator (diameter: 5 μm; threshold: 20 %), both performed in Arivis. All blobs within a region of width 100 μm, corresponding to the neural plate, were selected and their mean signal intensities were extracted. Such values were binned in 10 μm intervals, along the animal-vegetal axis, and plotted as a function of distance from the embryonic margin. The maximum of the binned values for each sample was used for normalization within that embryo.

### Gradient analysis using FCS

FCS measurements were carried out in the extracellular space at increasing distances from the embryonic margin, and the fluorescence count rates (CR) were recorded. The average CR corresponding to background autofluorescence was subtracted by taking FCS measurements in wildtype embryos. The CR values within a given sample were normalized to the maximum CR obtained for that sample. Resulting values from several such samples were binned in 40 μm intervals, averaged and plotted as a function of distance from the margin.

### smFISH foci analysis

Individual smFISH foci were segmented using the Blob finder analysis operator in Arivis. The blob radius was set to 1 μm, which was found to efficiently detect the most foci. A threshold < 5 was always applied but re-adjusted for each image manually to facilitate the detection of the maximum number of spots that could be identified visually. A rectangular region of interest, corresponding to the neural plate, was selected across the animal-vegetal axis and divided into 10 μm blocks using a custom-written Python script (Abbate, M., Technical support, Arivis). Blob counts within each 10 μm bins were recorded and plotted as a function of distance from the embryonic margin. For analysis of endogenous expression profiles, normalization was done within each sample to the maximum of the binned value for that sample. For analysis of *tbxta* expression in Heparinase I-injected samples, the average of the maxima for all control samples was used for normalization.

### Statistical test

In order to determine whether two datasets were significantly different from each other, a t-test was performed using GraphPad. It was assumed that the datasets follow a normal distribution and an un-paired t-test (with equal variance) was applied to compute the p-values. A p-value < 0.05 was considered as the measure for statistical significance.

### Probe sets for smFISH

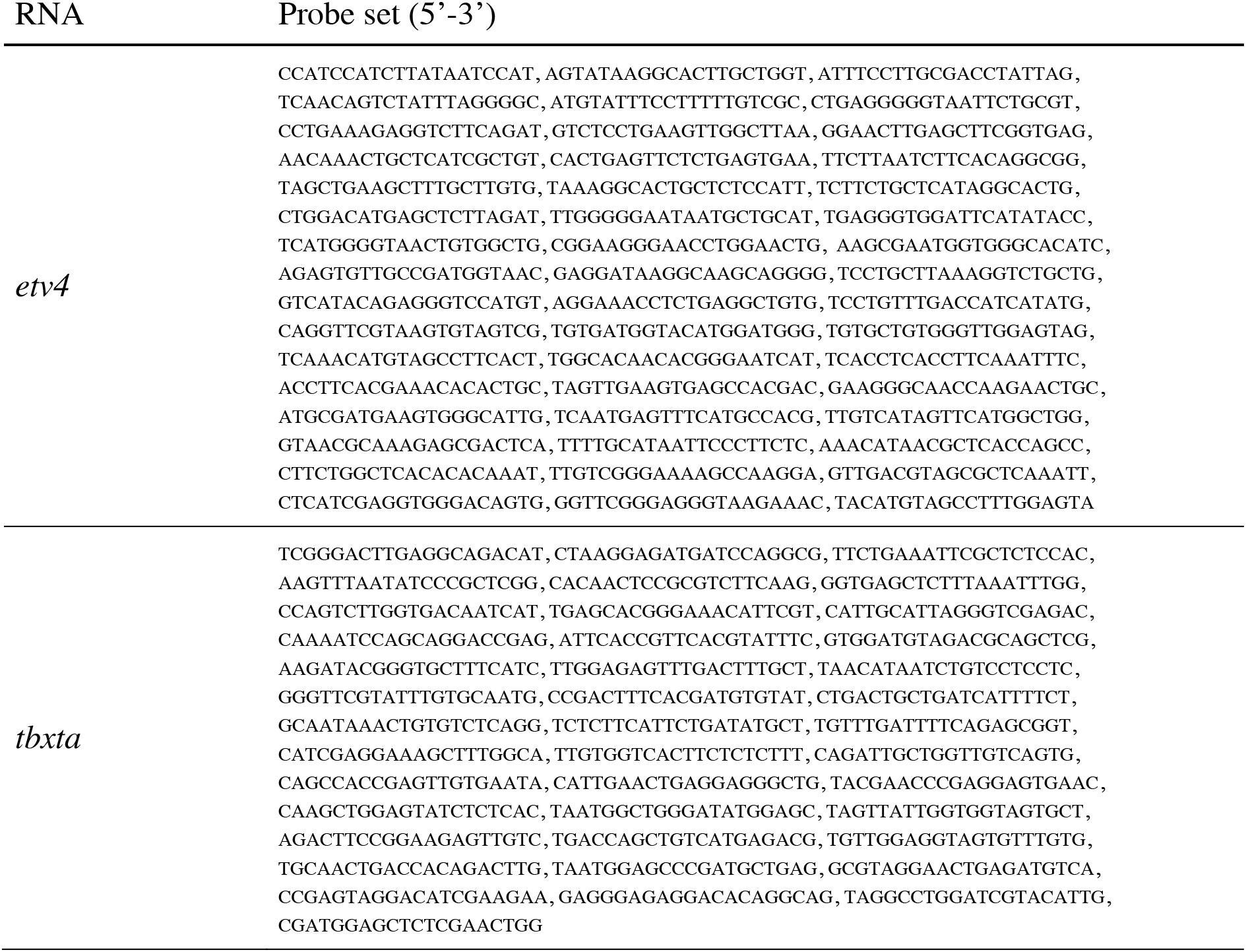

