## Supplementary figures and images for "Real-time monitoring of endogenous Fgf8a gradient attests to its role as a morphogen during zebrafish gastrulation"

### Supplementary figure 1

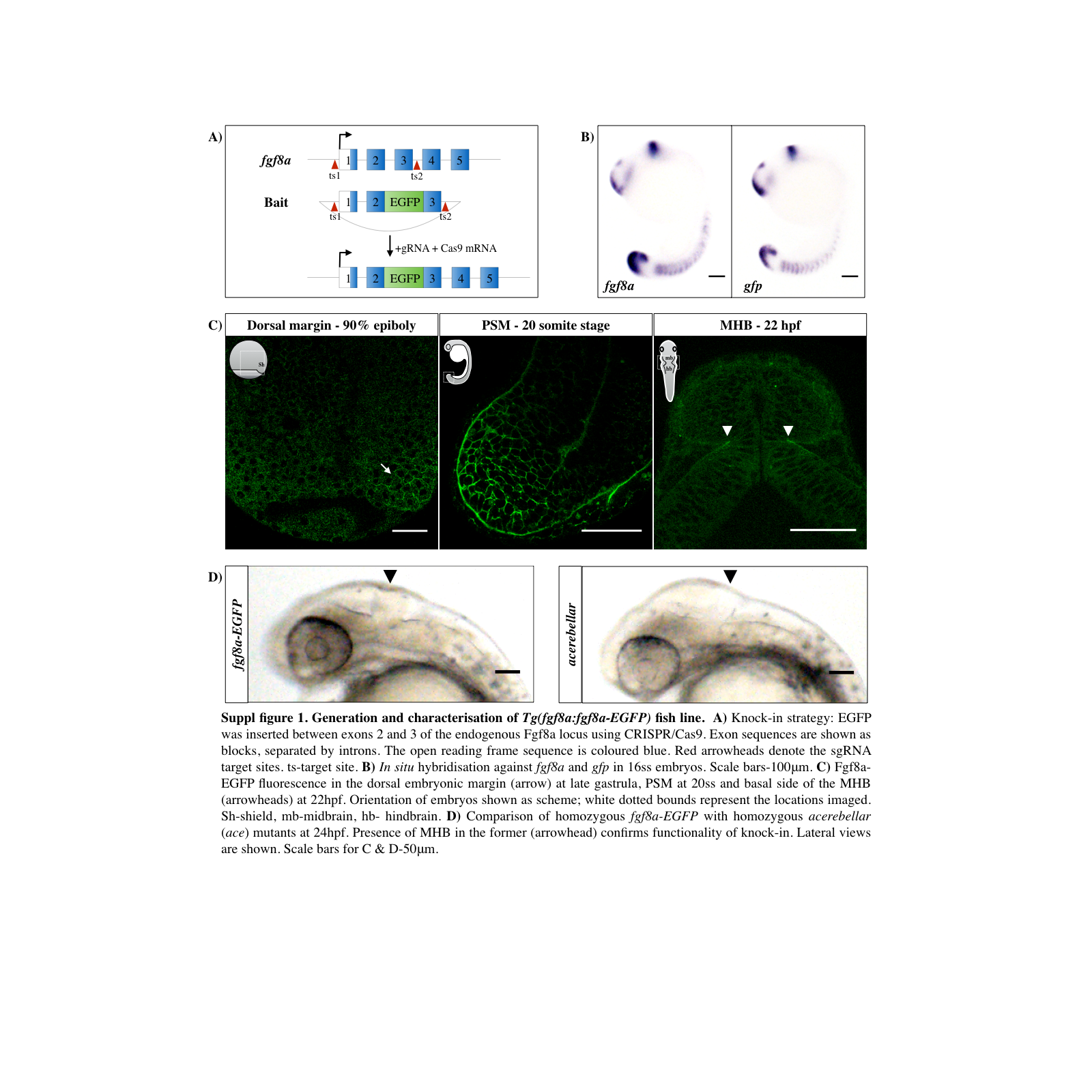

### Supplementary figure 2

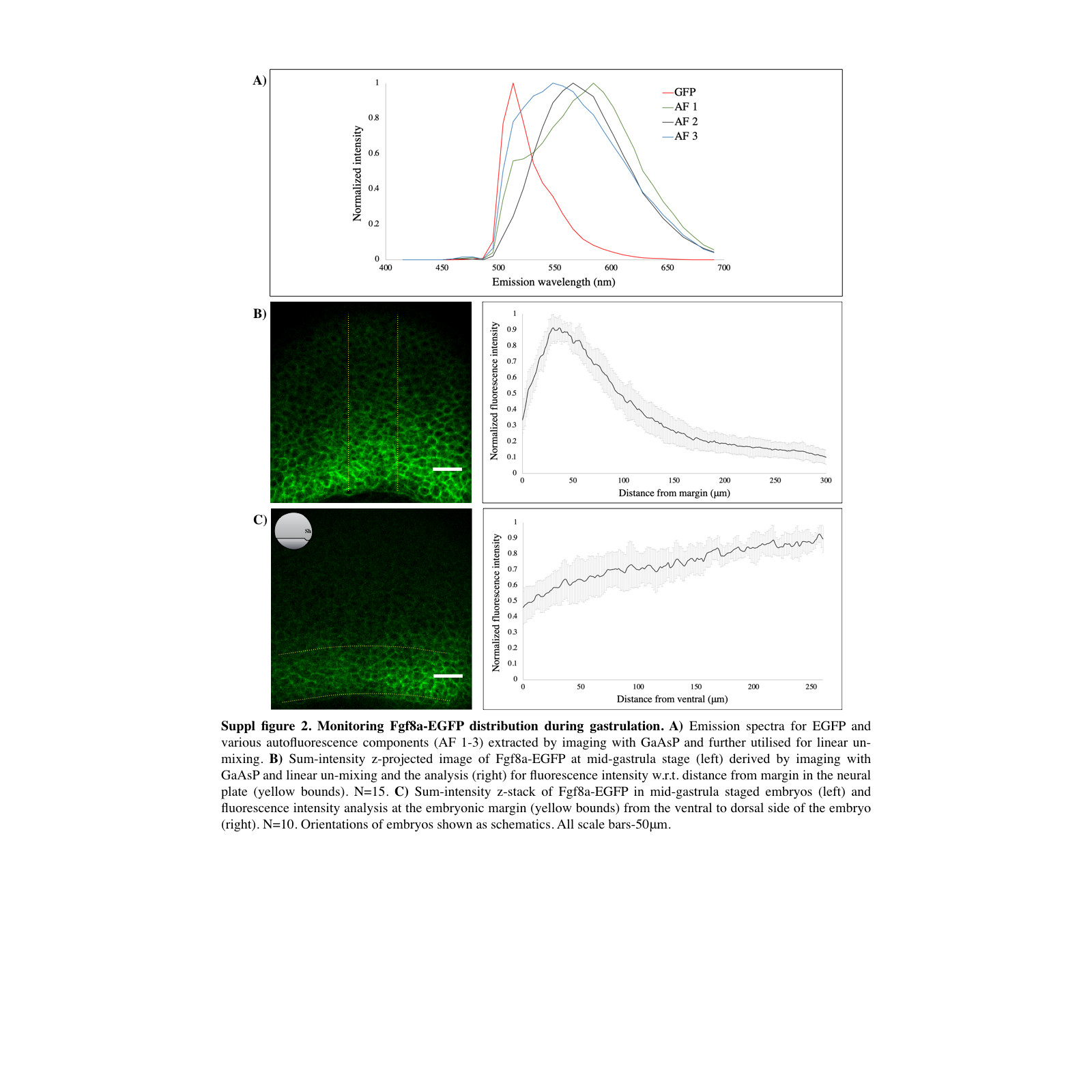

### Supplementary figure 3

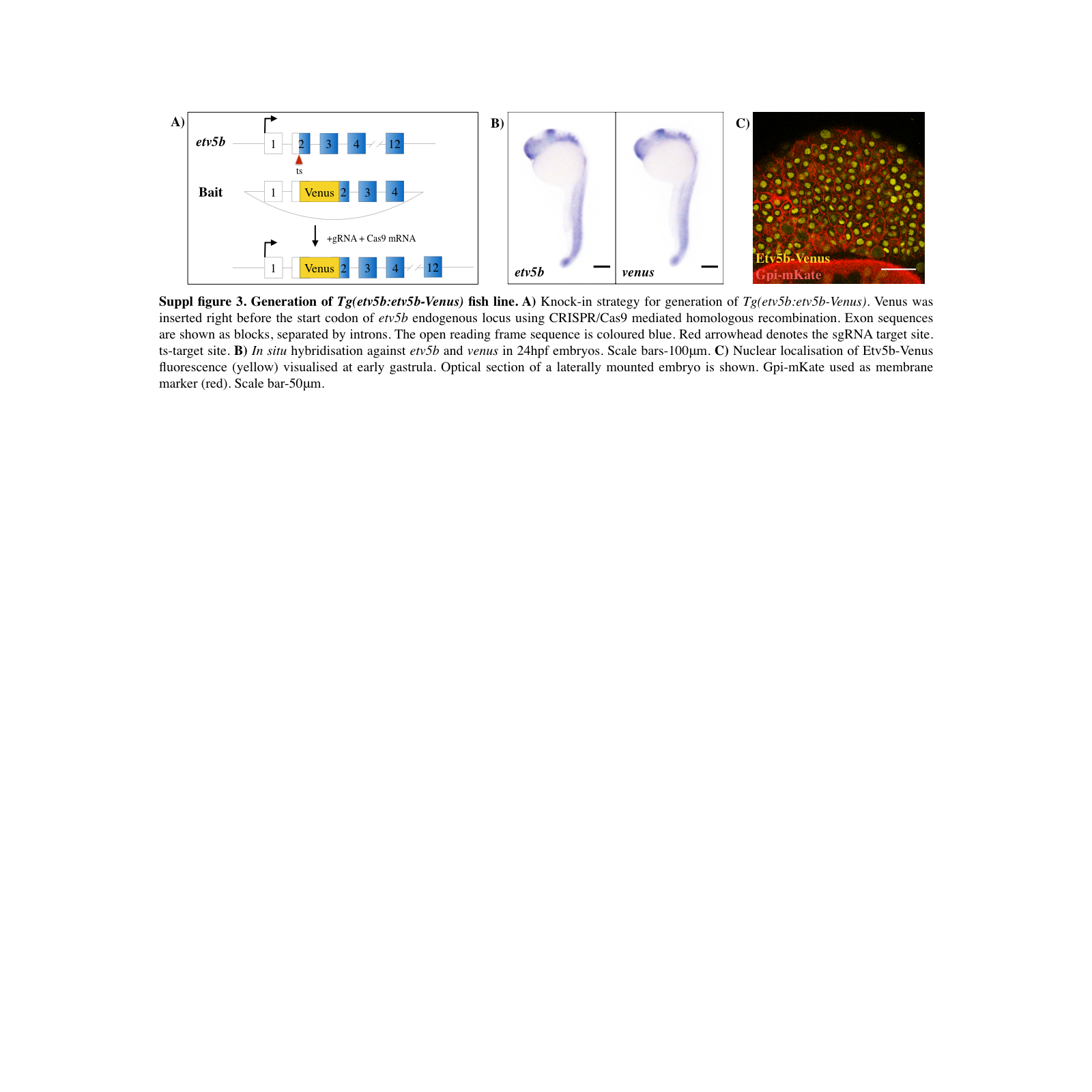

### Supplementary figure 4

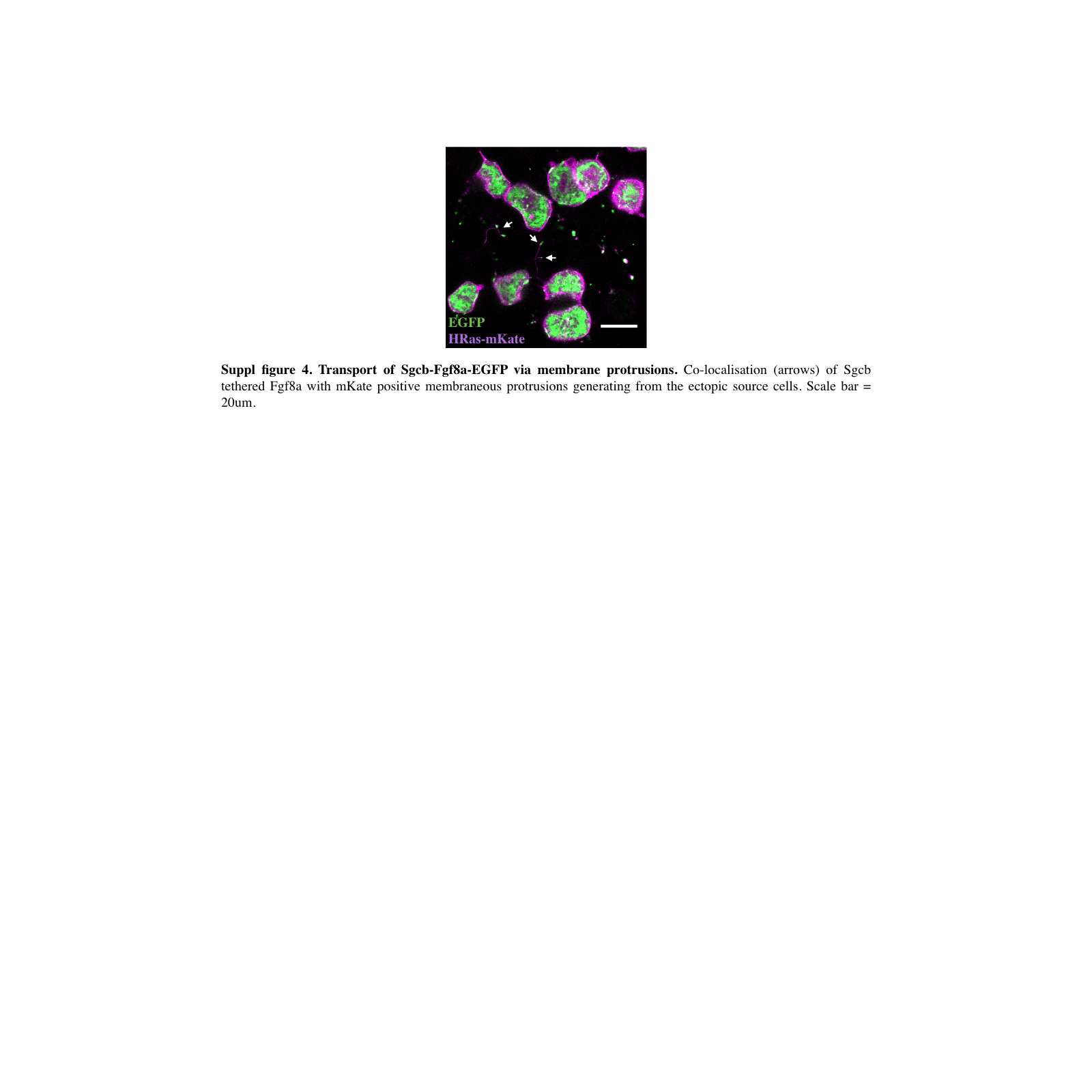
